# Reactivation of the X-linked *Nexmif* Gene Corrects Mosaic NEXMIF Deficiency in Heterozygous Female Mice

**DOI:** 10.1101/2025.11.29.691251

**Authors:** KathrynAnn Merth, Heng-Ye Man

## Abstract

We have previously demonstrated that heterozygous (HET) female mice lacking one copy of the X-linked gene *Nexmif* display autistic-like phenotypes, memory impairments, and deficits in synapse and neuron morphology. Due to random X Chromosome Inactivation (XCI), the HET mouse brain contains two populations of neurons: NEXMIF-expressing cells (wildtype, WT) and NEXMIF-lacking cells (knockout, KO). Interestingly, because KO cells contain a normal WT copy of *Nexmif* on the inactivated X chromosome (Xi), we wondered whether the silenced Xi-*Nexmif* could be reactivated to restore NEXMIF expression in neurons as a strategy to correct this mosaic deficiency in HET mice. To this end, we first tested pharmacological inhibition of XCI maintenance and found that intracortical administration of the DNA methylation inhibitor 5-aza-2’-deoxycytidine combined with resveratrol (Aza+Resveratrol) increased NEXMIF expression in HET mice. Using a gene-specific approach, we developed a *NEXMIF*-targeted CRISPR activation (CRISPRa) system and found that it selectively increases *NEXMIF* transcription in human female cells and *in vivo* in WT female mice with minimal off-target effects. Importantly, CRISPRa restored NEXMIF expression in the KO neurons of HET primary cultures, effectively correcting XCI-driven mosaicism. These findings demonstrate that pharmacological- and especially CRISPRa-mediated reactivation of the Xi may serve as a strategy for the reversal of neuronal and behavioral impairments in *Nexmif* HET conditions.

## Introduction

Individuals with autism spectrum disorder (ASD) display impairments in communication, social interaction deficits, and restricted/repetitive behaviors (1,2). Loss-of-function variants in the X-linked gene *NEXMIF* have been associated with NEXMIF syndrome, a developmental encephalopathy characterized by autistic features, intellectual disability (ID), and seizures in both male and female carriers (3,4). The NEXMIF protein is expressed specifically in neurons, localized primarily to the nucleus, and is widely distributed in the developing fetal brain. Findings from our studies and others revealed important roles for NEXMIF in early neural circuit formation, neurite outgrowth, and neuron migration (5–9). Our previous work demonstrates that in heterozygous (HET) female mice, *Nexmif* haploinsufficiency leads to morphological and synaptic impairments, including reduced dendritic spine density, attenuated dendritic branching and length, and downregulation of synaptic proteins (10). Furthermore, consistent with clinical observations in female patients (4), HET mice display autistic-like behaviors including altered social preference, repetitive behaviors, mild communication deficits, anxiety, memory impairment, and seizures (10). Given the critical role of NEXMIF in early neurodevelopment and the broad deficits arising from its loss, it is crucial to determine whether these cellular and behavioral impairments can be reversed.

In all female mammalian cells, the process of X chromosome inactivation (XCI) results in a unique genetic scenario in which only one X chromosome is active (Xa), while the other X is inactivated (Xi) to balance the gene dosage between the sexes (11). During XCI, the long noncoding RNA *Xist* accumulates on the future Xi, prompting the stable silencing of the entire chromosome (11,12). Recent studies have shown that *Xist* expression is regulated by proteins, i.e. XCI factors (XCIFs), which are normally involved in cell growth and proliferation, such as those in the BMP/TGF-β superfamily and PI3K/AKT pathway (13–16). These XCIFs, as well as *Xist* RNA binding proteins (RBPs), have been reported to promote the expression and/or localization of *Xist* to the Xi (13,17). XCI also involves epigenetic regulations such as the spreading of repressive H3K27me3 marks and methylation by the DNA methyltransferase 1 (DNMT1) (14,18–21).

While XCI ensures dosage compensation, it presents an interesting cellular scenario when a X-linked gene contains a loss-of-function variant, such as in heterozygous females with NEXMIF syndrome. We previously demonstrated that half of the neurons in HET female mice express NEXMIF and function as WT cells since the Xa contains the *Nexmif* allele, whereas the other half function as KO cells since the Xi contains the *Nexmif* allele, resulting in mosaic NEXMIF expression (10). Intriguingly, we found that although the WT cells in HET mouse brains express NEXMIF, they show impaired morphology and synaptic defects similar to the KO cells (10), demonstrating that loss of *Nexmif* in half of the neuronal population results in an overall dysregulation.

Studies suggest that the activity status of genes on the Xi chromosome can be reversed via X-chromosome reactivation (XCR). In primary neurons and mouse models of *Mecp2* Rett syndrome (16,17,22) and *Fmr1* Fragile X syndrome (23–26), XCR has been induced to restore the loss of these X-linked genes. Small molecule-mediated inhibition of XCIFs, *Xist* RBPs, histone deacetylation, and DNA methylation has shown promising Xi-linked gene reactivation, leading to rescue of neuronal deficits (16,22,27–30). However, these pharmacological approaches are often accompanied by off-target effects on other X-linked genes and even autosomal targets (31). Interestingly, recent innovations in CRISPR-based gene activation (CRISPRa) systems, which employ a catalytically-inactive Cas9 (dCas9) fused to transcriptional activators guided to the target gene by a promoter-specific RNA (32), provide a more target-specific and effective approach for Xi-gene reactivation. Notably, CRISPRa has been used to successfully reactivate the Xi-linked tumor suppressor gene *FOXP3* in human breast cancer cells (33), Xi-*MECP2* in human stem cell-derived Rett syndrome neurons (34), and in autosomal gene-based haploinsufficient disorders (35–37). These studies demonstrate the capability of CRISPRa in reversing key cellular deficits with no detectable off-target effects. However, whether CRISPRa or epigenetic modulation can be applied to reactivate Xi-*Nexmif* in the HET mouse brain remains unknown.

Here, we investigated the efficacy of epigenetic inhibitors and the CRISPRa system in increasing NEXMIF expression via reactivation of Xi-*Nexmif*. We found that inhibition of DNA methylation by 5-Aza-2’-deoxycytidine (Aza) combined with resveratrol resulted in an increase in NEXMIF expression in HET mouse neurons. Further, lentiviral delivery of a *NEXMIF*-specific CRISPRa system led to robust *NEXMIF* upregulation from the Xi in cultured human and mouse female neurons and *in vivo* in wild-type (WT) female mice. We confirmed that transduction of cultured HET neurons with the CRISPRa virus successfully induces *Nexmif* re-expression in the KO neurons. These findings establish the feasibility of pharmacological and CRISPR approaches in reactivating *Nexmif* from the Xi chromosome, suggesting their potential in rescuing neuronal impairments and autistic-like phenotypes in *Nexmif* HET female mice.

## RESULTS

### *Nexmif* XCR via Pharmacological Inhibition of XCI Factors (XCIFs) in Mouse and Human Neurons

Establishment and maintenance of XCI involves *Xist* upregulation by XCIFs and repressive DNA/histone modifications (13,20,38,39). Indeed, the pharmacological inhibition of these processes has shown favorable increases in target gene expression from the Xi (15,40,41). Thus, we wanted to explore the effects of pharmacological XCI inhibition on *Nexmif* expression in HET mice. Primary cortical neurons (DIV 3) prepared from P0 WT males, WT females, and HET female littermates were treated with either 0.1% DMSO control, the DNA methyltransferase 1 (DNMT1) inhibitor 5-aza-2’-deoxycytidine (Aza, 0.2μM) + Resveratrol (20μM), the JAK/STAT inhibitor AG-490 (3 μM), or the histone deacetylase inhibitor RG-2833 (5 μM) every 3 days until DIV 9 and collected for RT-qPCR at DIV 10 **(Fig. 1A**). Treatment with Aza+Resveratrol significantly increased mRNA levels of *Nexmif*, but not *Mecp2*, in HET and WT female cortical neurons (**Fig. 1B)**. Neither AG-490 nor RG-2833 produced significant changes in *Nexmif* or *Mecp2* expression. Moreover, Aza+Resveratrol did not alter *Nexmif* or *Mecp2* expression in WT male neurons (**Fig. 1B-C**), suggesting a saturating expression rate or a mechanism that limits the expression flexibility of genes on the active X.

**Figure 1.**
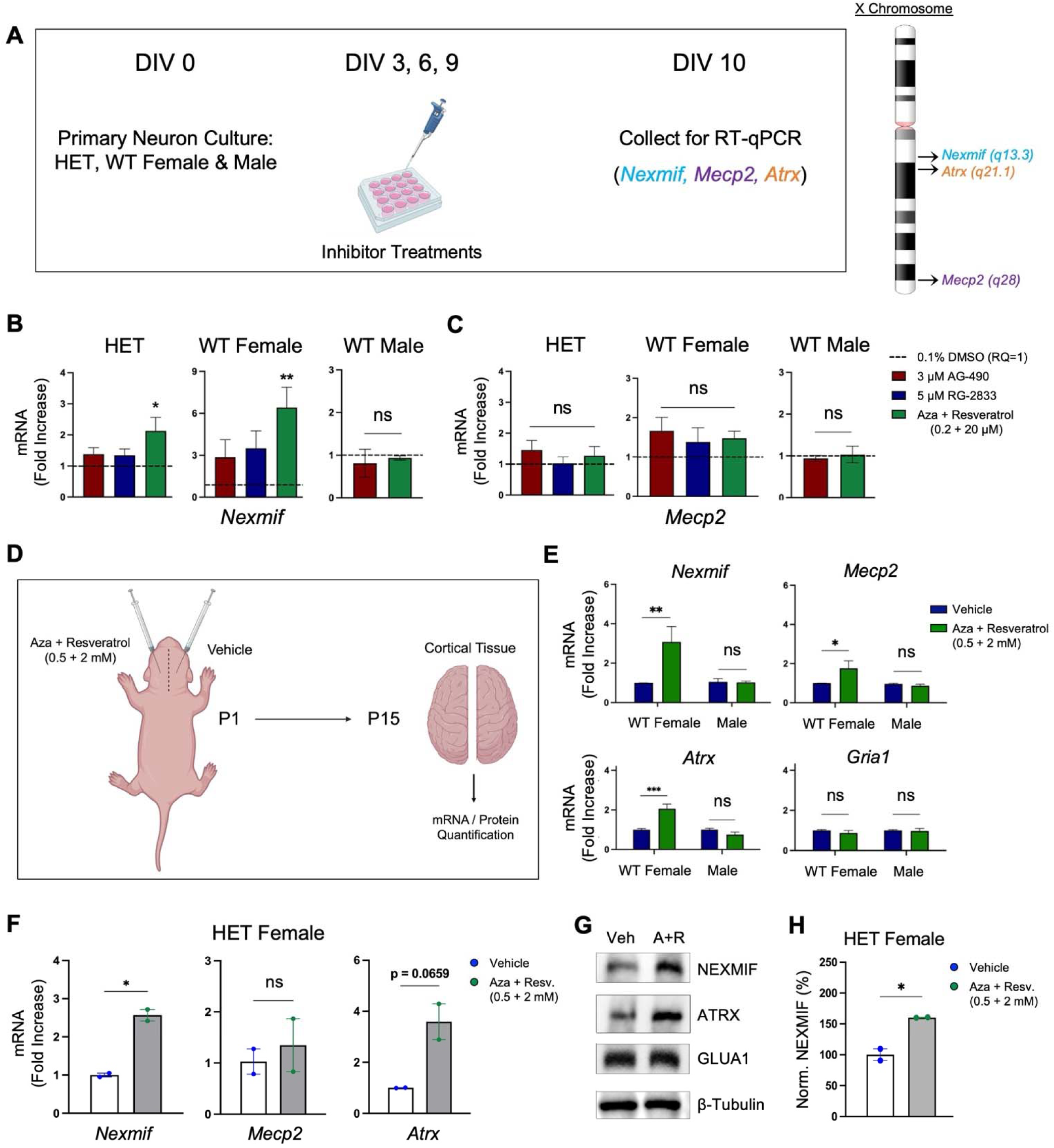
The DNA Methyltransferase 1 (DNMT1) inhibitor 5-aza-2’-deoxycytidine (Aza)+Resveratrol increases NEXMIF expression in WT and HET female mice. ***A*,** Cortical neurons were cultured from P0 WT male, WT female, and HET female mice and treated with 0.1% DMSO or X chromosome reactivation (XCR) compounds every 3 days until DIV 10, at which point mRNA was extracted for qPCR quantification. ***B-C*,** Aza+Resveratrol increased *Nexmif,* but not *Mecp2,* mRNA levels in HET and WT female neurons, and did not alter expression in WT male neurons (n=8 WT F, 8 HET, 5 WT M). Aza+Resveratrol (green bar, DNMT1 inhibitor); RG-2833 (blue bar, HDAC 1/3 inhibitor); AG-490 (red bar, JAK2 inhibitor). ***D*,** Vehicle or Aza+Resveratrol were injected unilaterally into the cortices of P1 WT male, WT female, and HET female mice, and cortical tissue was collected for mRNA and protein quantification at P15. ***E*,** Aza+Resveratrol significantly increased *Nexmif*, *Mecp2*, and *Atrx* mRNA levels in WT females, but not in WT males, by P15. mRNA levels of the autosomal gene *Gria1* were unaltered in both WT males and females (n=4F and 4M). ***F*,** Aza+Resveratrol increased *Nexmif* and *Atrx* mRNA, but not *Mecp2*, in HET females by P15. ***G-H*,** Western blot images (G) and quantification (H) showing increased NEXMIF protein levels in P15 HET females. Mean±SEM. One-way ANOVA with Bonferroni’s Multiple Comparisons (B-C) and Two-tailed t test (E,F,H). *p<0.05, **p<0.01, ***p<0.001; ns, not significant.

Using Aza+Resveratrol as our candidate inhibitor, we then performed unilateral intracortical brain injections in P1 WT males, WT females, and HET littermates: 3 μl of vehicle was injected into the right hemisphere and 3 μl of Aza+Resveratrol (0.5 mM + 2 mM) was injected into the left hemisphere of each mouse (**Fig. 1D**). At P15, cortical tissue from each hemisphere was collected for mRNA and protein quantification. Aza+Resveratrol treatment increased *Nexmif* mRNA and protein levels in the drug-treated hemisphere of WT females and of HETs (**Fig. 1E-H**), but did not alter *Nexmif* expression in WT males, relative to the vehicle-injected hemisphere. Aza+Resveratrol treatment also did not alter the expression of the autosomal gene *Gria1* (**Fig. 1E**), suggesting specificity for the X chromosome. However, Aza+Resveratrol also increased *Mecp2* and *Atrx* levels in WT female mice, and trended towards increased *Atrx* levels in HET mice (**Fig. 1E**), indicating reactivation of other genes aside from *Nexmif*.

To directly visualize *Nexmif* reactivation in the neurons of the brain, P1 HET mice received bilateral ventricular (ICV) brain injections of either vehicle or Aza+Resveratrol, followed by perfusion and brain slice staining at P30. Consistently, we found increased NEXMIF immunosignal in the medial prefrontal cortex (mPFC) (**Fig. 2A,C**). Moreover, this increase in NEXMIF expression was accompanied by a reduction in H3K27me3 methylation, indicating a weakened suppression in gene expression (**Fig. 2B-C**).

**Figure 2.**
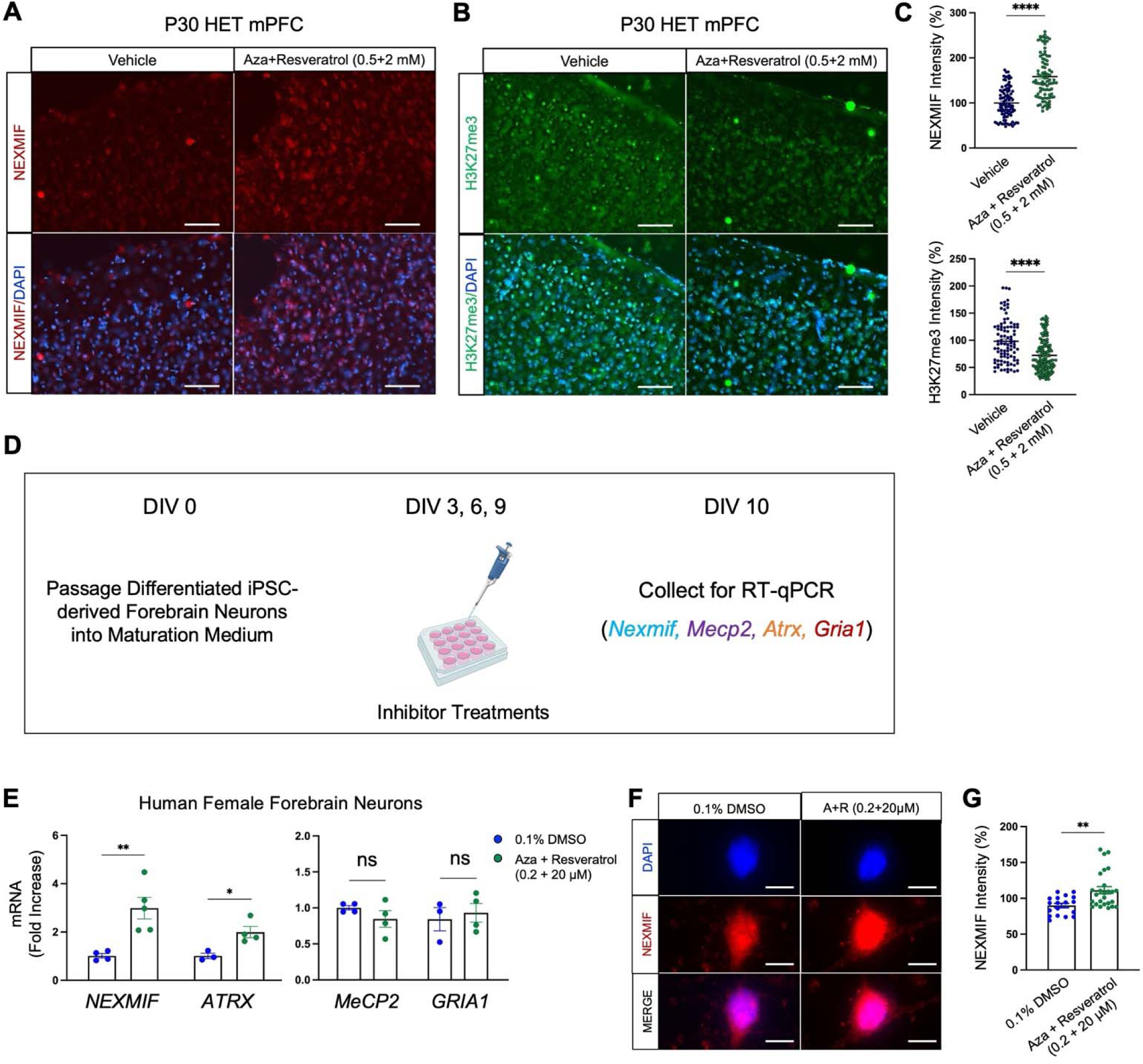
Aza+Resveratrol increases NEXMIF expression in the HET brain and in Human Female Neurons. ***A-B*,** Vehicle or Aza+Resveratrol (0.5+2μM) was bilaterally injected into the ventricles of P1 HET female mice. Brain tissue was collected at P30. Immunostaining of (A) NEXMIF and (B) H3K27me3 in the medial prefrontal cortex (mPFC) of injected HET mice. Scale bar=200μm. ***C*,** Quantification of nuclear intensity revealed a significant increase in NEXMIF expression (top graph) and a significant reduction in H3K27me3 expression (bottom graph) in Aza+Resveratrol-injected brains, relative to vehicle. ***D*,** Human female iPSC-derived neural progenitor cells were differentiated into forebrain neurons and passaged into maturation medium (DIV 0). At DIV 3, the forebrain neurons were treated with 0.1% DMSO control or Aza+Resveratrol every 3 days until DIV 10, at which point mRNA was extracted for RT-qPCR quantification. ***E*,** *NEXMIF* and *ATRX* mRNA levels were increased, but not *MECP2* or *GRIA1*. ***F*,** Representative immunostaining images of DAPI (top row) and NEXMIF (middle row) from DMSO-treated (left panel) and Aza+Resveratrol-treated (right panel) human female forebrain neurons at DIV 10; Scale bar=5μm. ***G*,** Quantification of (F) revealed a significant increase in NEXMIF immunosignal induced by Aza+Resveratrol (n=18 control and 26 treated cells). Mean±SEM. Two-tailed student’s t test (C,E,G). *p<0.05, **p<0.01, ****p<0.0001; ns, not significant.

To further determine the rescue effect in human cells, human female iPSC-derived neural progenitor cells were differentiated into forebrain neurons. Three days after the neurons were plated in maturation medium (DIV 3), the neurons were treated with either 0.1% DMSO or Aza (0.2 μM) + Resveratrol (20 μM) every 3 days until DIV 10, at which point the cells were either lysed for mRNA quantification or fixed for immunostaining (**Fig. 2D**). As expected, Aza+Resveratrol increased *NEXMIF* expression (**Fig. 2E**). To examine the specificity, we found that in addition to *NEXMIF*, the mRNA levels for *ATRX*, but not *MeCP2* or *GRIA1*, were upregulated by the treatment (**Fig. 2E**). Consistently, NEXMIF protein expression was also significantly increased in the Aza+Resveratrol-treated neurons as indicated by the elevated immunosignal (**Fig. 2F-G**). These results demonstrate that DNA methylation inhibition by Aza+Resveratrol can effectively stimulate *NEXMIF* gene transcription.

### CRISPRa–Induced *NEXMIF* Transcription in Human Female Cells

While the pharmacological approach shows effectiveness in reactivating *Nexmif* expression, it also regulates the expression of additional X-linked genes. In seeking a more specific XCR approach, we utilized the CRISPR/dCas9-VP64 (CRISPRa) system and constructed a guide RNA (gRNA) sequence specific to the human *NEXMIF* promoter. The dCas9-VP64 plasmid was co-transfected with either control-gRNA or NEX-gRNA in human female-derived HEK cells for 48 hr, followed by mRNA and protein quantification. The CRISPRa system increased *NEXMIF* mRNA and protein levels in HEK cells without altering *MeCP2* or *ATRX* expression, indicating specificity for *NEXMIF* (**Fig. 3A-C**). We then tested the CRISPRa system in human female iPSC-derived forebrain neurons. Upon plating in maturation medium (DIV 0), the forebrain neurons were co-transduced with the dCas9-VP64 lentivirus together with either control- or NEX-gRNA lentivirus. 7 d later, immunostaining of the neurons revealed a robust increase in NEXMIF expression by CRISPRa relative to control (**Fig. 3D-E**), demonstrating the efficacy of CRISPRa in enhancing *NEXMIF* gene transcription.

**Figure 3.**
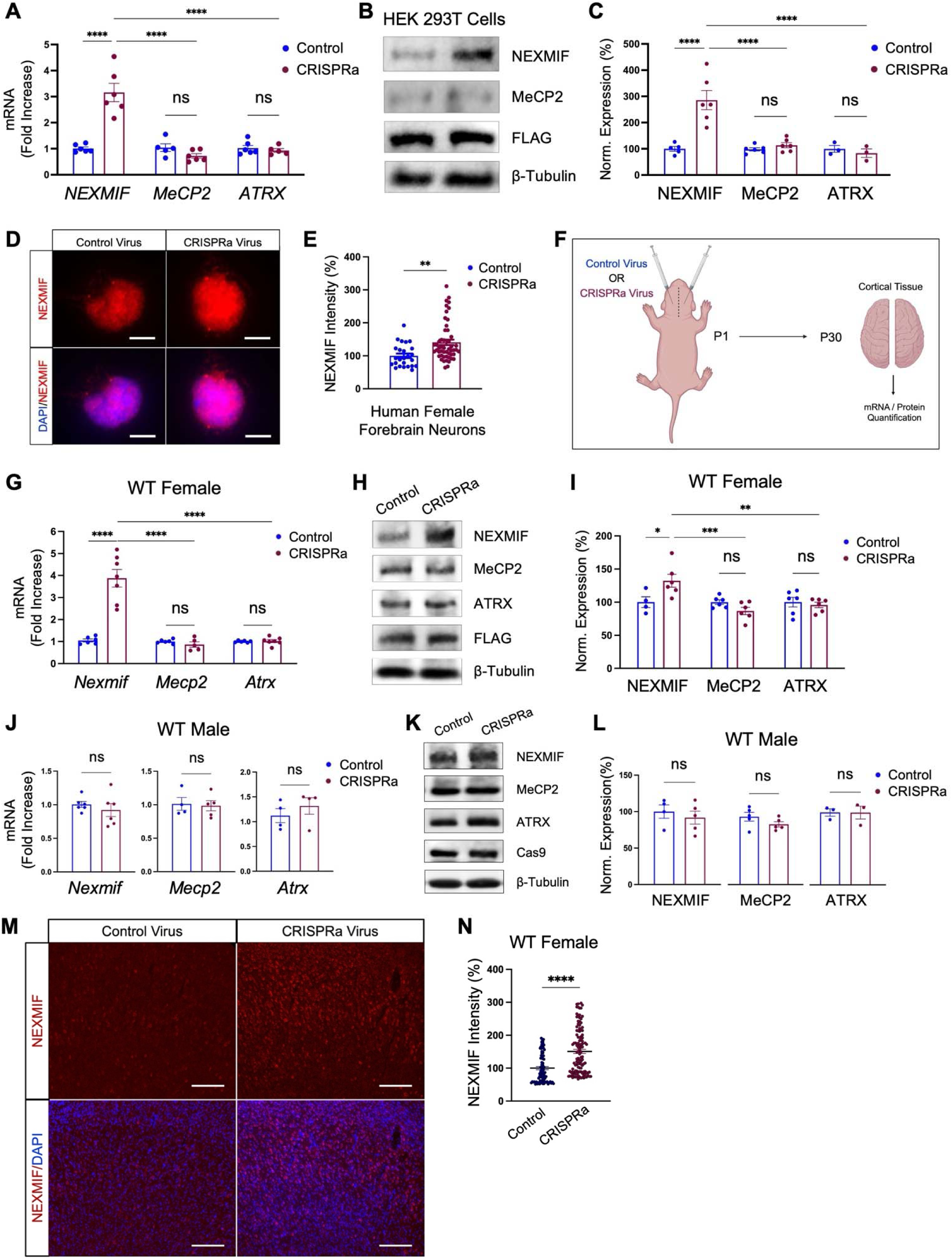
CRISPRa targeting of the *NEXMIF* promoter increases NEXMIF expression in Human Female Cells and WT Female Mice. ***A-C*,** Validation of the CRISPRa system: HEK293T cells were co-transfected with dCas9-VP64 + Control-gRNA (blue bar) or NEX-gRNA (purple bar) for 48h. CRISPRa led to increased *NEXMIF* mRNA (A) and protein (B-C), with no changes in *MeCP2* or *ATRX* expression (N=5-6). FLAG indicates the presence of dCas9-VP64. ***D*,** Human female forebrain neurons (DIV 3) were treated with dCas9-VP64 lentivirus together with control- or NEX-gRNA lentivirus. Representative images of treated neurons (DIV 10) immunostained against NEXMIF (top) merged with DAPI (bottom). Scale bar=5µm. ***E*,** Quantification of (D) revealed that CRISPRa significantly increased NEXMIF nuclear expression. n=20-40 neurons/group. ***F,*** Control or CRISPRa lentivirus was bilaterally injected into the ventricles of P1 WT female and male mice. Cortical brain tissue was collected at P30 for mRNA and protein quantification. ***G-I,*** CRISPRa significantly and specifically increased *Nexmif* mRNA (G) and protein expression (H-I), with no changes in *Mecp2* or *Atrx* expression. ***J-L,*** In WT male mice, CRISPRa did not significantly alter *Nexmif, Mecp2,* or *Atrx* mRNA (J) or protein expression (K-L). ***M,*** NEXMIF immunostaining in the medial prefrontal cortex (mPFC) of P30 WT female mice injected with either control or CRISPRa lentivirus at P1. Scale bar=400μm. ***N,*** Quantification of NEXMIF immunostaining revealed significantly increased nuclear expression in cortical neurons in response to specific activation of the *Nexmif* promoter by CRISPRa, relative to that of control-injected mice. Mean±SEM. Two-way ANOVA with Sidak’s Post Hoc Test (A-C, G-I) or a Two-tailed student’s t test (E,J,L,N). *p<0.05; **p<0.01; ***p<0.001; ****p<0.0001; ns: non-significant.

### Lentiviral CRISPRa Increases NEXMIF in WT Female Mice

We then validated the effectiveness of the CRISPRa virus *in vivo* in the brains of WT female mice. Following ICV brain injection of control or CRISPRa virus into WT female and male mice at P1, cortical brain tissue was collected at P30 for mRNA/protein quantification and brain slice imaging (**Fig. 3F**). We found that the CRISPRa virus significantly increased *Nexmif* mRNA and protein expression in WT female mice relative to controls, with little to no change in the expression of *Mecp2* or *Atrx* (**Fig. 3G-I**). Consistently, brain slice staining also revealed increased NEXMIF immunosignal in the mPFC of CRISPRa-injected WT female mice, relative to controls (**Fig. 3M-N**). In contrast, WT male mice injected with CRISPRa showed little to no alterations in *Nexmif*, *Mecp2*, or *Atrx* expression, relative to control male mice (**Fig. 3J-L**). These results validate the efficacy and specificity of CRISPRa for *Nexmif* upregulation *in vivo* in the living female mouse brain, as well as suggests that *Nexmif* reactivation occurs at the Xi given the lack of effect in males.

### CRISPRa Induces Xi-*Nexmif* Reactivation in HET Neurons

We have shown that CRISPRa can effectively increase NEXMIF expression in cultured human female cells and in living WT female mice. To determine whether the CRISPRa system indeed induces XCR of Xi-*Nexmif* in the KO cells of HET mice, we cultured cortical neurons from P0 WT males, WT females, and HET female littermates. At DIV 0, the neurons were transduced with either control or CRISPRa virus and left to incubate until DIV 7, at which point the neurons were fixed for immunostaining and quantification of the percentage of NEXMIF-expressing (NEXMIF+) cells (**Fig. 4A**). As expected, the total and average NEXMIF intensities in the HET + Control condition were significantly reduced relative to those of the WT female + Control condition (**Fig. 4B-D**), and the ratio of NEXMIF+ to NEXMIF-cells in the HET + Control condition was ∼55:45 (**Fig. 4E-F**). However, in the HET + CRISPRa condition, we found that the percentage of NEXMIF+ cells was significantly increased, accompanied by an increase in NEXMIF intensity (**Fig. 4B-F**), indicating that the CRISPRa system can sufficiently reactivate Xi-*Nexmif* in the KO cells of HET mice. In contrast, CRISPRa treatment did not alter MECP2 intensity in WT or HET female neurons (**Fig. 4G-H**), supporting the specificity of the CRISPRa system towards the *Nexmif* protomer. Furthermore, CRISPRa did not alter NEXMIF or MECP2 intensities in WT male neurons (**Fig. 4I-K**), which is in line with reactivation of the *Nexmif* gene at the Xi rather than general facilitation of NEXMIF expression from the active X.

**Figure 4.**
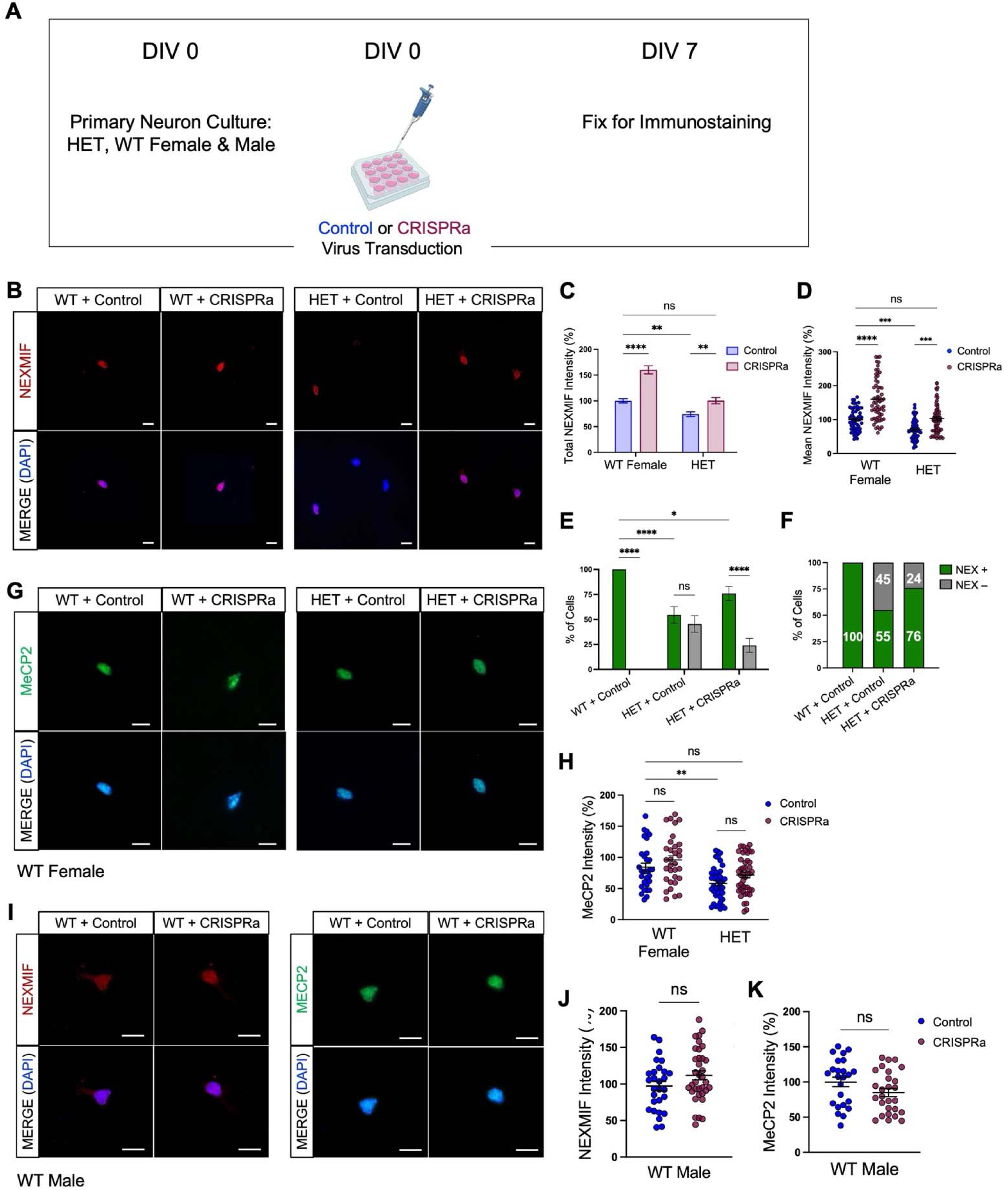
CRISPRa induces Xi-*Nexmif* Reactivation in Primary HET Neurons. ***A*,** Cortical neurons were cultured from P0 WT male, WT female, and HET female mice and transduced with either control or CRISPRa lentivirus from DIV 0➔7, at which point the cells were fixed for NEXMIF and MECP2 immunostaining. ***B*,** Representative images of transduced WT and HET female neurons (DIV 7) immunostained against NEXMIF (top) merged with DAPI (bottom). Scale bar=15µm. ***C-D*,** Quantification of total (B) and average (C) NEXMIF intensity revealed that CRISPRa transduction significantly increased nuclear expression in both WT female and HET neurons, relative to that of control (N = 62 neurons/condition). ***E-F*,** CRISPRa increased the percentage of NEXMIF+ (NEX+) neurons in HET culture, indicating Xi-Nexmif reactivation in NEXMIF- (NEX-) neurons (N = 62-74 neurons/condition). ***G*,** Representative images of transduced WT and HET female neurons immunostained against MECP2 (top) merged with DAPI (bottom). Scale bar=15µm. ***H,*** CRISPRa did not significantly alter MECP2 nuclear intensity in WT female or HET neurons (N = 39-40 neurons/condition.) ***I*,** Representative images of transduced WT male neurons immunostained against NEXMIF (top left panels) or MECP2 (top right panels) merged with DAPI (bottom panels). Scale bar=15µm. ***J-K,*** In WT male neurons, CRISPRa treatment did not significantly alter neither NEXMIF (J) nor MECP2 (K) nuclear intensity (J: N = 28-35 neurons/condition; K: N = 24-27 neurons/condition). Mean±SEM. *p<0.05, **p<0.01, ***p<0.001, ****p<0.0001; ns, not significant. Two-way ANOVA with Sidak’s Multiple Comparisons (C-E,H) and Two-tailed t test (J-K). N = 2 WT Females, 2 HETs, and 2 WT Males.

## DISCUSSION

In this study, we examined the feasibility and therapeutic promise of Xi-*Nexmif* reactivation strategies for the alleviation of neuronal and behavioral defects in HET female mice. Our findings revealed that inhibition of DNA methylation via Aza+Resveratrol increases *Nexmif* expression in WT and HET female mice, but also results in increases in *Mecp2* and *Atrx*. By contrast, the *NEXMIF* promoter-specific CRISPR activation approach enabled precise reactivation of Xi-*NEXMIF* in human female cells and in WT and HET female mice. Specifically, CRISPRa increased the number of NEXMIF-expressing cells in HET cultures to almost match that of WT female cultures, with little to no alterations in WT male neurons. These findings underscore the specificity and efficacy of CRISPRa-mediated gene reactivation, and suggest the potential of the CRISPRa system to improve neuronal and behavioral impairments *in vivo* in HET mice.

Epigenetic pharmacological inhibitors, although strong in gene reactivation (16,17,28,42), lead to global alterations in chromatin states and thus nonspecific gene dysregulation (43). Despite this, with further clinical testing, Aza+Resveratrol may hold translational promise given that Aza, also known as Decitabine, is currently used for the treatment of acute myelogenous leukemia via its activation of silenced tumor repressor genes (44), and clinical trials have shown that Resveratrol is well-tolerated (45). Nonetheless, the clinical utility of epigenetic inhibitors will ultimately depend on strategies capable of directing their effects to specific genes. In contrast, the CRISPRa system is advantageous over epigenetic inhibition due to its gene-specific transcriptional enhancement, resulting in low off-target effects. Here, the successful reactivation of Xi-*Nexmif* in the NEXMIF-lacking neurons indicates that, at least in cultured neurons, XCI-induced mosaicism can be corrected with specificity and without the use of XCI inhibitors.

Since the WT neurons of HET mice still express NEXMIF, viral expression of CRISPRa may cause NEXMIF overdosage in these neurons, potentially leading to functional dysregulation. However, CRISPRa has been shown to induce only modest upregulation of active genes, especially depending on the strength of the transcriptional activator type (e.g., VP64 vs. VP128) (46). Indeed, since NEXMIF expression was unaltered by CRISPRa in WT males, whose neurons contain only Xa-*Nexmif*, it is possible that CRISPRa only stimulates *Nexmif* at the Xi, adding an extra layer of specificity. While our work exhibited high specificity for *Nexmif* relative to *Mecp2* and *Atrx*, there is still a potential for off-target gene activation if the gRNA used shares partial homology with other promoter regions. Nevertheless, continued rigorous testing and validation of various *Nexmif* gRNAs should mitigate this issue (47). Future studies using CRISPRa in HET mouse brains will employ dCas9-chromatin immunoprecipitation (ChIP) sequencing to obtain a more comprehensive map of any off-target binding that may occur.

Future work will also confirm whether brain injection of viral CRISPRa in HET mice indeed leads to increased NEXMIF expression *in vivo*, and if so, full or partial rescue of the cellular and behavioral impairments is expected. In addition, the minimum level of NEXMIF re-expression needed to obtain optimal reversal of HET phenotypes remains unclear. Such knowledge will allow us to establish the optimal effective viral dosage of CRISPRa for the rescue of NEXMIF syndrome-related deficits. Overall, our findings provide a foundation for the development of therapeutics for NEXMIF syndrome and other X-linked haploinsufficient disorders.

## MATERIALS & METHODS

### Animal care and use

Mouse colonies were maintained in the Laboratory Animal Science Center (LASC) at the Boston University Charles River Campus on a C57BL/6J genetic background. *Nexmif* HET female mice (X^+^X^-^) were crossed with WT male mice to obtain WT male, WT female, and HET female littermates for experimental use. Following all *in vivo* injection experiments, IACUC-approved euthanasia methods were used to sacrifice mice prior to tissue collection, specifically via cryo-anesthesia for neonates or CO_2_ inhalation at a flow rate of 4 L/min in the home-cage for 3 min followed by cervical dislocation for adolescent mice.

### Genotyping

DNA was isolated from tail snips using the Hot Shot Method. A single tail snip was collected from each mouse at either postnatal (P) day 0–1 (prior to injections or primary culture), or at the time of weaning on P21, and placed in an Eppendorf tube. Tail snips were incubated for 30min at 95°C in 75µl of alkaline lysis buffer (25mM NaOH, 0.2M EDTA). The tubes were allowed to cool at room temperature (RT) for 5min before 75µl of neutralization buffer (40mM Tris-HCl, pH 5) was added. The tubes were mixed by vortexing and 1µl of DNA extract was used in the following PCR protocol run on a Biorad DNA engine Tetrad 2 Peltier Thermal Cycler: 94°C for 5min followed by 30 cycles of 98°C for 15s, 55°C (WT) or 59°C (KO) for 30s and 72°C for 60s. The resulting PCR fragments were run on a 1.2% agarose gel with ethidium bromide to label and visualize DNA under ultraviolet light. Two sets of primers were used to genotype *Nexmif* HET mice. One set (WT) was targeted against the exon 4 region using the primers: 5’−aggacttgcttaggttgcttcatggaa−3’ and 5’−cttaaattgctctacctcaagaccacca−3’ with an expected PCR fragment of 949bp. The other set (KO) was targeted against the KO cassette using the primers: 5’−cacacctccccctgaacctgaaag−3’ and 5’−cccacgaagggatcataccctgta−3’ with an expected PCR fragment of 794bp.

### Primary Neuron Culture

Cortical brain tissue was dissected from P0–1 *Nexmif* HET mice and WT littermate controls and used for primary neuron culture. The sex and genotype of each mouse were determined via PCR prior to culturing (described above). Each brain region was individually digested in a digestion buffer [papain (15mg/ml in Hanks balanced salt solution, Sigma-Aldrich #4762), L-Cysteine (4mg/ml in Hanks balanced salt solution, Sigma-Aldrich #C7352), and 0.5M EDTA pH 7.0] for 20min at 37°C, then triturated in a trituration buffer [0.1% DNase (Thermo Fisher #PA5-22017), 1% ovomucoid (Sigma-Aldrich #T2011)/1% bovine serum albumin (Sigma-Aldrich #05470) in Dulbecco’s modified Eagle’s medium (DMEM)] to fully dissociate neurons. Dissociated neurons were counted and plated on 18-mm circular coverslips (Carolina #633013) in 60-mm Petri dishes (five coverslips/dish) and 6-well culture plates that had been coated in poly-l-lysine (Sigma-Aldrich #P2636; 100μg/ml in borate buffer) overnight at 37°C then washed three times with sterile deionized water and left in plating medium [minimal essential medium (500mL) containing 10% fetal bovine serum (Atlanta Biologicals #S11550), 5% horse serum (Atlanta Biologicals #S12150), 31mg/ml L-cysteine, 1% penicillin/streptomycin (Corning #30-002-Cl), and 1% L-glutamine (Corning #25-005-Cl) before cell plating. The day after plating, plating medium was replaced by feeding medium (Neurobasal medium supplemented with 1% horse serum, 2% SM1, and 1% penicillin/streptomycin and 1% L-glutamine) which was supplemented with 5′-fluoro-2′-deoxyuridine (10μm; Sigma-Aldrich #F0503) after 4d *in vitro* to suppress glial growth.

### Human iPSC-Derived Neural Progenitor Cell and Forebrain Neuron Culture

Coverslips and 6-well plates were coated with poly-L-ornithine (Sigma #P4957) followed by laminin (Sigma #L2020) to support neural differentiation. Human female iPSC-derived neural progenitor cells (NPCs, Stem Cell Technologies #200-0620) were thawed and plated in STEMdiff™ Neural Progenitor Medium (#05833) for initial expansion for 7 d. NPCs were then passaged and plated in STEMdiff™ Forebrain Neuron Differentiation Medium (#08600) for another 7 d. The differentiated cells were then plated onto coverslips (2x10^5^) or 6-well plates (8x10^5^) and maintained in STEMdiff™ Forebrain Neuron Maturation Medium (#08605). The medium was replaced every 3 d until completion of experiments.

### Preparation and Use of Small-Molecule Inhibitors

All inhibitors used for primary neuron culture were dissolved in DMSO (Sigma Aldrich #D8418). The inhibitors used were 3µM AG-490 (JAK2 inhibitor, Tocris #0414), 5µM RG-2833 (HDAC1/3 inhibitor, Selleck Chemicals #S7292), or the DNA methyltransferase 1 inhibitor combination of 0.2µM 5-Aza-2’-deoxycytidine (Aza, Sigma-Aldrich #A3656) + 20µM Resveratrol (Selleck Chemicals #S1396). Cultured neurons (DIV 3) were treated with either 0.1% DMSO control or the inhibitors every 3 days until DIV 9 and lysates were collected for mRNA quantification at DIV 10. For *in vivo* brain injections, Aza and Resveratrol were resuspended together in vehicle (0.9% NaCl, 0.5% methylcellulose, 4.5% DMSO in sterile deionized H_2_O) at final concentrations of 0.5mM and 2mM, respectively. The solution was aliquoted and stored at - 20°C until use.

### Plasmids

The lentiviral (LV) plasmid pLV hUbC-dCas9 VP64-T2A-GFP (dCas9-VP64) was a gift from Charles Gersbach (Addgene #53192). The LV empty guide RNA (gRNA) control plasmid was a gift from Bejamin Ebert (Addgene #57822). The human *NEXMIF* promoter-specific LV gRNA was custom ordered from Millipore Sigma (#LV03) using the sequence CTTAAGTTAAGAGTAGGTAA.

### Plasmid Transfection

For HEK293T cell transfections, cells were plated 1:10 in 60-mm culture dishes. Two days later, cells were co-transfected using the target DNA plasmids and polyethylenimine reagent (PEI, Polysciences #23966). For one dish of cells, the plasmids and PEI were first separately diluted in 50µl of serum-free DMEM and then mixed and incubated at RT for 20min to form the transfection complex. The complex was then added dropwise to the cells and left to incubate for 48h prior to lysate collection.

### Virus Preparation

LVs were produced by co-transfecting HEK293T cells with the DNA plasmids and viral packaging and envelope proteins (pRSV/REV, pMDLg/pRRE, and pCMV-VSV-G) using PEI. Sodium pyruvate (Thermo Fisher #11360070) was added to the medium 24h later to supplement the cells. Conditioned medium containing the viral particles was harvested 48h later and filtered through a 0.45-μm filter. PEG-it Virus Precipitation Solution (System Biosciences #LV810A) was added to the medium, and the mixture was left to incubate at 4°C for 72h. The mixture was centrifuged at 1500g for 30min at 4°C and the viral pellet was resuspended in sterile 1X phosphate buffered saline (PBS). The virus was divided into 10μl aliquots and stored at −80°C. The LV packaging constructs were gifts from Didier Trono (Addgene #12251 and #12253) and Bob Weinberg (Addgene #8454).

### Neuron Transduction

For viral transductions, human female forebrain neurons or neurons cultured from WT male, WT female, or HET female littermates were plated on coverslips and infected with LV. 24h later, the medium was replaced with fresh medium, and the cells were incubated at 37°C in a 5% CO_2_ incubator for the desired amount of time.

### Mouse Brain Injections

P1 WT male, WT female, or HET female littermates were cryo-anesthetized on wet ice for 3min prior to unilateral (intracortical) or bilateral intracerebroventricular (ICV) injection of LVs or drugs on a chilled stage using a 10-μl syringe with a sterile 32-gauge needle (Hamilton #7653-01). Fast Green FCF dye (1μl, Thermo Scientific #A16520-14) was added to the LV aliquots to visualize and confirm successful injection. Following injection, pups were warmed on an isothermal heating pad with home-cage bedding before being returned to the dam.

### Immunocytochemistry (ICC) of Cultured Neurons

Neurons were fixed for 8min in a 4% paraformaldehyde (PFA) / 1X PBS at RT. Cells were rinsed twice in 1X PBS followed by membrane permeabilization for 10min in 0.3% Triton-X-100 (Sigma Aldrich #T8787) in 1X PBS. Cells were then rinsed twice in 1X PBS followed by incubation with primary antibodies overnight at 4°C, washed three times with cold 1X PBS, and incubated with Alexa Fluor-conjugated fluorescent secondary antibodies (1:500) for 1h at RT. Cells were then washed three times with cold 1X PBS, with the first wash containing Hoechst (1:10,000, Thermo Fisher #62249) and mounted to microscopy glass slides with Prolong Gold antifade mounting reagent (Thermo Fisher #P36930) for subsequent visualization. Mounted coverslips were kept overnight in the dark at RT before imaging. For ICC, primary and secondary antibodies were diluted in IHC-Tek™ Antibody Diluent pH 7.4 (IHCWorld, #IW-1000), which does not require the serum blocking step.

### Immunohistochemistry (IHC) of Brain Slices

P30 WT or HET female mice underwent transcardial perfusion with ice cold 1X PBS followed by 4% PFA prior to collection of the brain. Brains were post-fixed in 4% PFA for 4h and cryoprotected in 30% sucrose/1X PBS for 48h. Brains were then placed in molds to be rapidly frozen in Tissue-Tek OCT (Sakura #4583) with dry ice and stored at −80°C until they were cut into 30μm sections using a LEICA CM1850 cryostat (LEICA Biosystems). Sections were mounted onto SuperFrost microscope slides (Fisher Scientific #12-550-15) and stored at −20°C until staining. To prepare for immunostaining, sections were hydrated in 1X PBS for 30min followed by permeabilization in 1% Triton X-100/1X PBS for 1h. Sections were then incubated in primary antibodies overnight at 4°C in a humidity chamber, washed three times 10min each with cold 1X PBS, and incubated in Alexa Fluor-conjugated fluorescent secondary for 1h at RT. Brain slices were then washed three times with cold 1X PBS, with the first wash containing Hoechst (1:10,000) and mounted under a rectangular coverslip with Prolong Gold anti-fade mounting reagent. Slides were allowed to dry in the dark at RT overnight and stored at −20°C prior to subsequent visualization. For IHC, primary and secondary antibodies were diluted in IHC-Tek™ Antibody Diluent, which does not require the serum blocking step.

### Western Blot

Brains from P15 or P30 mice were dissected on ice immediately after sacrifice. For a ∼30mg piece of cortical tissue, or a 60 mm HEK cell dish, ∼75μl of ice-cold lysis buffer [50mM Tris-HCl pH 8, 150mM NaCl, 1% Triton X-100, 0.5% sodium deoxycholate (SDOC), 0.1% sodium dodecyl sulfate (SDS), supplemented with 100X protease inhibitor cocktail (Apex Bio #K1011)] was added and samples were homogenized followed by sonication (for 10s). Samples were then centrifuged for 15min at 13,000rpm at 4°C in a microcentrifuge. The tubes were placed on ice, and the supernatant was carefully aspirated and placed into a pre-cooled tube. Samples were subjected to a BCA assay according to the manufacturer’s protocol (Thermo Fisher #23225) to determine protein concentrations. Protein levels were normalized with the lysis buffer, and an equal volume of 2X sample reducing buffer [5% SDS, 150mM Tris-HCl pH 6.8, 0.05% Bromophenol Blue (Sigma-Aldrich #B0126), 5% fresh 2-mercaptoethanol (Sigma-Aldrich #M3148)] was added to the samples. The lysates were then boiled for 10min at 95°C. SDS-PAGE was performed to separate proteins of interest using standard procedures. Samples were run on 6–12% gels at 110V for 1hr. Proteins were transferred at 150mA overnight to PVDF membranes and blocked for 1h in 5% bovine serum albumin (BSA, Sigma Aldrich #A2153) prepared in 1X tris-buffered saline supplemented with 0.1% Tween (TBST). After blocking, membranes were probed with the appropriate primary antibody diluted in Signal Enhancer Hikari Buffer (Nacalai USA #NU00101) overnight at 4°C. Membranes were washed 3x 5min each in 1X TBST and then incubated with the appropriate secondary antibody for 1h. After secondary incubation, membranes were washed 3x with 1X TBST. Blots were visualized using the Azure Radiance Plus chemiluminescence detection system (Azure Biosystems #AC2103) on the Sapphire Biomolecular Imager (Azure Biosystems) and analyzed using NIH Fiji (ImageJ, RRID:SCR_002285).

### Antibodies

Primary antibodies to the following proteins were used: rabbit anti-KIAA2022 [1:100 (brain slice for IHC) and 1:300 (cultured neurons for ICC), Sigma #HPA000407], rabbit anti-KIAA2022 (1:750 for WB, Biorbyt Orb312213), rabbit anti-β-tubulin III (1:1000 for WB, Sigma-Aldrich T2200), rabbit anti-GluA1 (1:1000 for WB, Homemade), rabbit anti-FLAG (1:1000 for WB, Cell Signaling #14793), rabbit anti-ATRX (1:1000 for WB, Cell Signaling #10321), mouse-anti MECP2 [1:1000 (WB) and 1:100 (ICC), Santa Cruz sc-137070), and rabbit-anti H3K27me3 (1:300 for IHC, Active Motif #39158). The following secondary antibodies were used: IgG-HRP for WB [1:10000; Thermo Fisher, mouse (#62-6520) and rabbit (#31460)], Alexa Fluor 555 [1:500, Thermo Fisher, rabbit (#A-21428)], and Alexa Fluor 488 [1:500, Thermo Fisher, mouse (#A-11001), rabbit (#A-11008)] for ICC and IHC.

### Reverse Transcription and Quantitative Polymerase Chain Reaction (RT-qPCR)

Total RNA was extracted from ∼30mg of mouse cortical tissue, 5x10^5^ primary mouse neurons, 8x10^5^ human female forebrain neurons, or a 60-mm HEK cell dish using the RNeasy Mini kit (Qiagen #74106). RNA concentrations were diluted with nuclease-free water (NEB #B1500S) to 1μg (brain tissue) or 500ng (cultured cells), followed by reverse transcription to cDNA using the EasyQuick RT MasterMix (CoWin Biosciences #CW2019). 1μl of the cDNA was added to the HotStart™ Universal 2X SYBR Green qPCR Master Mix (ApexBio Technology #K1170) in a 20µl volume and real-time fluorescence quantitative PCR was performed using the 7900HT Fast Real-Time PCR System (Applied Biosystems) to detect mRNA levels with the appropriate primers (**Supplementary Table 1**). Target gene expression was normalized to *Gapdh* prior to using the delta delta Ct method to determine the fold change in gene expression.

### Microscopy

Exposure time for the fluorescence signals was adjusted manually so the signals were within a full dynamic range. Once the parameters were set, they were fixed and used throughout image acquisition for each experiment.

**For ICC:** Fluorescent images were collected with a 40x and 63x oil-objectives on a ZEISS Axio Imager Z2 Upright Microscope using the ZEISS ZEN software.

**For IHC:** Brain sections were imaged with 20x air-objectives using a DS-Fi2 Color Camera on a Nikon Eclipse NiE using NIS-Elements software.

### Statistical Analysis

**For ICC, IHC, and Western Blot:** A One-Way ANOVA with Tukey’s or Bonferroni’s Post Hoc Test was used to determine significant differences in fluorescent intensity and mRNA expression between inhibitor treatments. Either a two-tailed student’s t test or a Two-Way ANOVA with Sidak’s Post Hoc Test was used, where appropriate, to determine significant differences in mRNA, protein expression, or fluorescent intensity between DMSO/Vehicle/Control- and Aza+Resveratrol/CRISPRa-treated neurons/brains.

All data are expressed as Mean±SEM (standard error of the mean) and were analyzed using GraphPad Prism 10 statistical software (GraphPad Software, Boston, MA). Sample sizes for neuronal and behavioral experiments were based on a power of 0.8 and an α=0.05, with effect sizes estimated based on our previous studies. All data were treated as parametric for statistics, and equal variances were assumed when comparing means across multiple groups. Outliers were defined as individual data points with a value larger or smaller than 2 standard deviations from the mean. p<0.05 is considered statistically significant. *p* values are presented as p>0.05 (ns, not significant), \**p*<0.05, \*\**p*<0.01, \*\*\**p*<0.001, and \*\*\*\**p*<0.0001.

## ACKNOWLEDGEMENTS

We would like to thank the Man Lab members for their insightful input. This work was supported by R01 MH130600, R21 MH133014, and R21 MH134174. The authors declare no competing financial interests.

## AUTHOR CONTRIBUTIONS

KO: Data Curation, Formal Analysis, Investigation, Methodology, Validation, Visualization, Writing – Original Draft, Writing – Review & Editing, Conceptualization; HYM: Conceptualization, Data Curation, Funding Acquisition, Investigation, Methodology, Project Administration, Supervision, Validation, Visualization, Writing – Review & Editing.

## REFERENCES

1. Landa RJ. Diagnosis of autism spectrum disorders in the first 3 years of life. Nat Clin Pract Neurol. 2008 Mar;4(3):138–47.

2. CDC. Autism Spectrum Disorder (ASD). 2024 [cited 2024 Oct 31]. Signs and Symptoms of Autism Spectrum Disorder. Available from: https://www.cdc.gov/autism/signs-symptoms/index.html

3. About XLID98 - XLID98 Foundation [Internet]. [cited 2025 Aug 16]. Available from: https://www.xlid98.org/about-xlid98

4. Stamberger H, Hammer TB, Gardella E, Vlaskamp DRM, Bertelsen B, Mandelstam S, et al. NEXMIF encephalopathy: an X-linked disorder with male and female phenotypic patterns. Genet Med Off J Am Coll Med Genet. 2021 Feb;23(2):363–73.

5. Cantagrel V, Haddad MR, Ciofi P, Andrieu D, Lossi AM, Maldergem LV, et al. Spatiotemporal expression in mouse brain of Kiaa2022, a gene disrupted in two patients with severe mental retardation. Gene Expr Patterns. 2009 Sept;9(6):423–9.

6. Ishikawa T, Miyata S, Koyama Y, Yoshikawa K, Hattori T, Kumamoto N, et al. Transient expression of Xpn, an XLMR protein related to neurite extension, during brain development and participation in neurite outgrowth. Neuroscience. 2012 July;214:181–91.

7. Magome T, Hattori T, Taniguchi M, Ishikawa T, Miyata S, Yamada K, et al. XLMR protein related to neurite extension (Xpn/KIAA2022) regulates cell–cell and cell–matrix adhesion and migration. Neurochem Int. 2013 Nov;63(6):561–9.

8. Gilbert J, Man HY. The X-Linked Autism Protein KIAA2022/KIDLIA Regulates Neurite Outgrowth via N-Cadherin and δ-Catenin Signaling. eneuro. 2016 Sept;3(5):ENEURO.0238-16.2016.

9. Gilbert J, O’Connor M, Templet S, Moghaddam M, Di Via Ioschpe A, Sinclair A, et al. NEXMIF/KIDLIA Knock-out Mouse Demonstrates Autism-Like Behaviors, Memory Deficits, and Impairments in Synapse Formation and Function. J Neurosci. 2020 Jan 2;40(1):237–54.

10. O’Connor M, Qiao H, Odamah K, Cerdeira PC, Man HY. Heterozygous Nexmif female mice demonstrate mosaic NEXMIF expression, autism-like behaviors, and abnormalities in dendritic arborization and synaptogenesis. Heliyon. 2024 Jan 24;10(3):e24703.

11. Maxfield Boumil R, Lee JT. Forty years of decoding the silence in X-chromosome inactivation. Hum Mol Genet. 2001 Oct 1;10(20):2225–32.

12. Jacobson EC, Pandya-Jones A, Plath K. A lifelong duty: how *Xist* maintains the inactive X chromosome. Curr Opin Genet Dev. 2022 Aug 1;75:101927.

13. Bhatnagar S, Zhu X, Ou J, Lin L, Chamberlain L, Zhu LJ, et al. Genetic and pharmacological reactivation of the mammalian inactive X chromosome. Proc Natl Acad Sci. 2014 Sept 2;111(35):12591–8.

14. Kumari D, Usdin K. Sustained expression of FMR1 mRNA from reactivated fragile X syndrome alleles after treatment with small molecules that prevent trimethylation of H3K27. Hum Mol Genet. 2016 Sept 1;25(17):3689–98.

15. Carrette LLG, Wang CY, Wei C, Press W, Ma W, Kelleher RJ, et al. A mixed modality approach towards Xi reactivation for Rett syndrome and other X-linked disorders. Proc Natl Acad Sci [Internet]. 2018 Jan 23 [cited 2024 Apr 19];115(4). Available from: https://pnas.org/doi/full/10.1073/pnas.1715124115

16. Przanowski P, Wasko U, Zheng Z, Yu J, Sherman R, Zhu LJ, et al. Pharmacological reactivation of inactive X-linked Mecp2 in cerebral cortical neurons of living mice. Proc Natl Acad Sci. 2018 July 31;115(31):7991–6.

17. Sripathy S, Leko V, Adrianse RL, Loe T, Foss EJ, Dalrymple E, et al. Screen for reactivation of MeCP2 on the inactive X chromosome identifies the BMP/TGF-β superfamily as a regulator of XIST expression. Proc Natl Acad Sci. 2017 Feb 14;114(7):1619–24.

18. Pandya-Jones A, Markaki Y, Serizay J, Chitiashvili T, Mancia Leon WR, Damianov A, et al. A protein assembly mediates Xist localization and gene silencing. Nature. 2020 Nov;587(7832):145–51.

19. Costanzi C, Pehrson JR. Histone macroH2A1 is concentrated in the inactive X chromosome of female mammals. Nature. 1998 June 11;393(6685):599–601.

20. Plath K, Fang J, Mlynarczyk-Evans SK, Cao R, Worringer KA, Wang H, et al. Role of histone H3 lysine 27 methylation in X inactivation. Science. 2003 Apr 4;300(5616):131–5.

21. Cotton AM, Price EM, Jones MJ, Balaton BP, Kobor MS, Brown CJ. Landscape of DNA methylation on the X chromosome reflects CpG density, functional chromatin state and X-chromosome inactivation. Hum Mol Genet. 2015 Mar 15;24(6):1528–39.

22. Carrette LLG, Wang CY, Wei C, Press W, Ma W, Kelleher RJ, et al. A mixed modality approach towards Xi reactivation for Rett syndrome and other X-linked disorders. Proc Natl Acad Sci [Internet]. 2018 Jan 23 [cited 2024 Apr 19];115(4). Available from: https://pnas.org/doi/full/10.1073/pnas.1715124115

23. Kumari D, Gazy I, Usdin K. Pharmacological Reactivation of the Silenced FMR1 Gene as a Targeted Therapeutic Approach for Fragile X Syndrome. Brain Sci. 2019 Feb 12;9(2):39.

24. Tabolacci E, Palumbo F, Nobile V, Neri G. Transcriptional Reactivation of the FMR1 Gene. A Possible Approach to the Treatment of the Fragile X Syndrome. Genes. 2016 Aug 17;7(8):49.

25. Haenfler JM, Skariah G, Rodriguez CM, Monteiro Da Rocha A, Parent JM, Smith GD, et al. Targeted Reactivation of FMR1 Transcription in Fragile X Syndrome Embryonic Stem Cells. Front Mol Neurosci. 2018 Aug 15;11:282.

26. Xie N, Gong H, Suhl JA, Chopra P, Wang T, Warren ST. Reactivation of FMR1 by CRISPR/Cas9-Mediated Deletion of the Expanded CGG-Repeat of the Fragile X Chromosome. Bardoni B, editor. PLOS ONE. 2016 Oct 21;11(10):e0165499.

27. Minkovsky A, Sahakyan A, Bonora G, Damoiseaux R, Dimitrova E, Rubbi L, et al. A high-throughput screen of inactive X chromosome reactivation identifies the enhancement of DNA demethylation by 5-aza-2′-dC upon inhibition of ribonucleotide reductase. Epigenetics Chromatin. 2015 Dec;8(1):42.

28. Lee HM, Kuijer MB, Ruiz Blanes N, Clark EP, Aita M, Galiano Arjona L, et al. A small-molecule screen reveals novel modulators of MeCP2 and X-chromosome inactivation maintenance. J Neurodev Disord. 2020 Nov 10;12(1):29.

29. Janiszewski A, Talon I, Chappell J, Collombet S, Song J, De Geest N, et al. Dynamic reversal of random X-Chromosome inactivation during iPSC reprogramming. Genome Res. 2019 Oct;29(10):1659–72.

30. Lou S, DJiake Tihagam R, Wasko UN, Equbal Z, Venkatesan S, Braczyk K, et al. Targeting microRNA-dependent control of X chromosome inactivation improves the Rett Syndrome phenotype. Nat Commun. 2025 July 4;16(1):6169.

31. Grimm NB, Lee JT. Selective Xi reactivation and alternative methods to restore MECP2 function in Rett syndrome. Trends Genet TIG. 2022 Sept;38(9):920–43.

32. Perez-Pinera P, Kocak DD, Vockley CM, Adler AF, Kabadi AM, Polstein LR, et al. RNA-guided gene activation by CRISPR-Cas9–based transcription factors. Nat Methods. 2013 Oct;10(10):973–6.

33. Cui X, Zhang C, Xu Z, Wang S, Li X, Stringer-Reasor E, et al. Dual CRISPR interference and activation for targeted reactivation of X-linked endogenous FOXP3 in human breast cancer cells. Mol Cancer. 2022 Feb 7;21:38.

34. Qian J, Guan X, Xie B, Xu C, Niu J, Tang X, et al. Multiplex epigenome editing of MECP2 to rescue Rett syndrome neurons. Sci Transl Med. 2023 Jan 18;15(679):eadd4666.

35. Tamura S, Nelson AD, Spratt PWE, Kyoung H, Zhou X, Li Z, et al. CRISPR activation rescues abnormalities in SCN2A haploinsufficiency-associated autism spectrum disorder [Internet]. bioRxiv; 2022 [cited 2025 Aug 8]. p. 2022.03.30.486483. Available from: https://www.biorxiv.org/content/10.1101/2022.03.30.486483v1

36. Jaudon F, Thalhammer A, Zentilin L, Cingolani LA. CRISPR-mediated activation of autism gene Itgb3 restores cortical network excitability via mGluR5 signaling. Mol Ther Nucleic Acids. 2022 Sept 13;29:462–80.

37. Colasante G, Qiu Y, Massimino L, Di Berardino C, Cornford JH, Snowball A, et al. In vivo CRISPRa decreases seizures and rescues cognitive deficits in a rodent model of epilepsy. Brain. 2020 Mar;143(3):891–905.

38. Jeon Y, Sarma K, Lee JT. New and Xisting Regulatory Mechanisms of X Chromosome Inactivation. Curr Opin Genet Dev. 2012 Apr;22(2):62–71.

39. Yang T, Ou J, Yildirim E. Xist exerts gene-specific silencing during XCI maintenance and impacts lineage-specific cell differentiation and proliferation during hematopoiesis. Nat Commun. 2022 Aug 1;13:4464.

40. Talon I, Janiszewski A, Chappell J, Vanheer L, Pasque V. Recent Advances in Understanding the Reversal of Gene Silencing During X Chromosome Reactivation. Front Cell Dev Biol [Internet]. 2019 Sept 3 [cited 2025 Sept 1];7. Available from: https://www.frontiersin.org/journals/cell-and-developmental-biology/articles/10.3389/fcell.2019.00169/full

41. Spaziano A, Cantone I. X-chromosome reactivation: a concise review. Biochem Soc Trans. 2021 Dec 17;49(6):2797–805.

42. Grimm NB, Lee JT. Selective Xi Reactivation and Alternative Methods to Restore MECP2 Function in Rett Syndrome. Trends Genet TIG. 2022 Sept;38(9):920–43.

43. Verma M, Banerjee HN. Epigenetic inhibitors. Methods Mol Biol Clifton NJ. 2015;1238:469–85.

44. Christman JK. 5-Azacytidine and 5-aza-2′-deoxycytidine as inhibitors of DNA methylation: mechanistic studies and their implications for cancer therapy. Oncogene. 2002 Aug;21(35):5483–95.

45. Berman AY, Motechin RA, Wiesenfeld MY, Holz MK. The therapeutic potential of resveratrol: a review of clinical trials. Npj Precis Oncol. 2017 Sept 25;1(1):35.

46. Becirovic E. Maybe you can turn me on: CRISPRa-based strategies for therapeutic applications. Cell Mol Life Sci CMLS. 2022 Feb 12;79(2):130.

47. Horlbeck MA, Witkowsky LB, Guglielmi B, Replogle JM, Gilbert LA, Villalta JE, et al. Nucleosomes impede Cas9 access to DNA in vivo and in vitro. eLife. 5:e12677.

